# Rapid sequence-based screening of structure-disrupting protein mutations

**DOI:** 10.64898/2026.02.24.707693

**Authors:** Jaemin Oh, Xiaoning Qian, Byung-Jun Yoon

## Abstract

Recent advances in AI-based protein structure prediction have dramatically reduced the cost of obtaining three-dimensional protein models and have become integral to modern protein engineering workflows. However, full structure prediction remains computationally prohibitive in high-throughput settings for mutation-based protein engineering, where thousands of candidate variants may need to be evaluated. In many such scenarios, the primary objective is not to resolve the complete structure of a candidate mutant, but rather to identify whether the introduced mutations are likely to induce substantial structural changes for rapid down-selection of candidates that conserve the wildtype structure. Protein language models (PLMs), trained solely on unlabeled natural protein sequence data, are known to encode rich structural information within their hidden representations. Motivated by this observation, we investigate a range of sequence-based ranking metrics derived from PLMs as efficient surrogates for structural deformation prediction. Through systematic evaluation across multiple proteins, mutation regimes, and structure-prediction backbones, we show that embedding distance—particularly the *L*^1^ distance between representations—provides a robust and computationally efficient signal for identifying structure-disrupting mutations. Our results demonstrate that sequence-level screening can substantially reduce the need for expensive structure prediction while preserving sensitivity to large structural perturbations, thereby providing the means to significantly speed up mutation-based protein design.

## Introduction

The tertiary structure of a protein is a primary determinant of its biochemical and biophysical function^1–3^. Fine-grained structural features govern molecular recognition events such as enzyme–substrate binding, receptor activation, and antigen–antibody interactions. For instance, neutralizing antibodies recognize viral antigens through precisely organized epitope–paratope contacts; even subtle geometric perturbations in these regions can diminish binding and facilitate immune escape. Modern protein engineering, therefore, aims to maintain essential structural and functional properties while optimizing other characteristics such as stability, expression, or affinity^4,5^. However, even single-point substitutions can induce substantial conformational re-arrangements that compromise protein function.

The risk of such conformational changes necessitates verifying the structures of engineered proteins before downstream experimental studies or clinical applications. X-ray crystallography remains the gold standard for resolving macromolecular structures, but its time-consuming and labor-intensive process presents a significant bottleneck in engineering workflows. Recent advances in AI-based structure prediction—most notably AlphaFold2 (AF2)^6^—have achieved near-experimental accuracy at dramatically reduced computational cost. Although there is ongoing debate regarding the extent to which AF2 predictions reflect mutational effects on stability or phenotype^7–9^, AF2 has clearly provided valuable structural insights and accelerated iterative protein design.

Beyond *de novo* generation, two broad classes of strategies dominate computational protein engineering. The first generates candidate amino-acid sequences and subsequently evaluates their predicted structures^10^. The second performs backbone-conditioned sequence design, in which a target structure is fixed and the model searches for compatible sequences^11,12^. The first approach incorporates structural information only at the end of the design process, whereas the second constrains exploration by enforcing structural compatibility from the outset. Incorporating structural information throughout the design process—for example, by constraining mutations whose predicted RMSD lies within a specified range—would enable more flexible and informed navigation of sequence space. However, running full structure prediction for every potential variant remains computationally prohibitive, particularly in high-throughput settings.

In this work, we address the problem of rapidly predicting whether a mutation is likely to induce a large structural change without performing full 3D structure prediction for each variant. This challenge is especially acute because a protein of length *L* admits 19*L* single-point mutants, rendering exhaustive structural evaluation infeasible. Our key insight is that modern protein language models, such as the Evolutionary Scale Modeling (ESM) family^13–15^, encode rich residue–residue interaction information that can be extracted in the form of predicted contact probabilities^16^. These contact-informed representations provide a computationally lightweight surrogate for structural reasoning: changes in protein language model outputs correlate with structural deviations in three-dimensional space.

Building on this observation, we systematically examine correlations between RMSD and ESM-derived scores across several proteins. We find that ESM-based contact-difference metrics and likelihood-based scores exhibit modest but statistically significant correlations with structural deformation. This result suggests that such representations can serve as efficient proxies for screening mutations in high-throughput design workflows, while also highlighting the broader utility of emergent structural signals in large protein language models.

## Background

### Evolutionary scale modeling

Evolutionary Scale Modeling (ESM) refers to a family of large protein language models trained on massive collections of natural amino acid sequences. ESM models adopt the masked language modeling objective: given a sequence

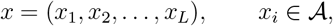

where 𝒜 is the amino acid alphabet. Then we consider *x*\_*ℳ*_, where a random subset ℳ ⊂1, …, *L* of positions is replaced by a special token. The model is then trained to predict the original residues at the masked positions by minimizing the negative log-likelihood

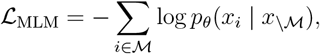

where *p*_*θ*_ denotes the probability distribution parameterized by the model. Because the model receives only sequence-level information, it must infer underlying patterns in natural amino acid sequences, many of which arise from structural and evolutionary constraints. Rives et al. ^13^ demonstrated that as ESM scales in depth and parameter count, hidden representations of ESM can be useful for many downstream tasks, including predictions of secondary structure, mutational effects, and residue-residue contacts. This emergent structural signal enables the use of ESM not only for masked-token inference but also as a foundation for downstream structure-prediction architectures such as ESMFold^14^.

### Multi-head attention

ESM adopts a multi-layer Transformer architecture^17^, whose core operation is *multi-head self-attention*. For sequences of length *L*, let

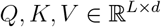

denote the query, key, and value representations, respectively. The scaled dot-product attention mechanism is

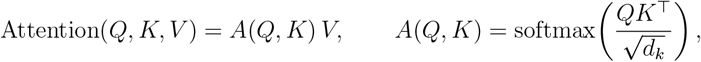

where the softmax is applied row-wise and *d*_*k*_ is the key dimension.

Each attention head *i* ∈ {1, …, *H*} is parameterized by projection matrices

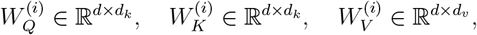

and produces

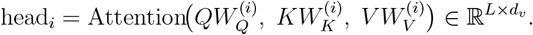

The outputs of the *H* heads are concatenated and projected:

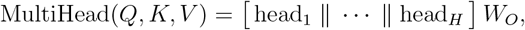

where 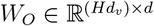. For convenience, the model typically sets *d*_*k*_ = *d*_*v*_ = *d/H*.

In the (ℓ + 1)-th Transformer layer, multi-head *self*-attention is applied to the layer input *h* ^(ℓ)^ ∈ ℝ^L × d:^

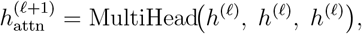

followed by the feed-forward sublayer and residual connections (omitted here for brevity). The initial representation *h*^(0)^ is obtained from the input amino acid sequence *x* ∈ {1, …, 20} ^L^ via a learned embedding,

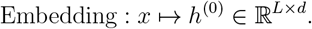

We omit additional Transformer components (layer normalization, positional encodings, feed-forward blocks) as they are not required for understanding the method introduced below. We refer the reader to the original references for full architectural details.

### Residue-residue contact predictions

A core hypothesis underlying the ESM family of protein language models is that, given sufficiently large training corpora and model capacity, three-dimensional structural information emerges implicitly within the model’s internal representations^13^. To test this hypothesis, Rao et al. ^16^ trained a logistic regression classifier in which attention-derived features serve as covariates and binary residue–residue contact indicators 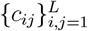 serve as response variables. Here *c*_*ij*_ = 1 if residues *i* and *j* are spatially proximal (based on C_*α*_ distance) and *c*_*ij*_ = 0 otherwise. Let *A*^(*l,h*)^ ∈ R^*L×L*^ denote the attention matrix^1^ from layer *l* ∈ {1, …, *N*_ℓ_} and head *h* ∈ {1, …, *N*_*h*_}. The model takes the form

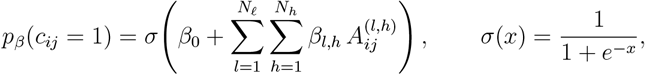

where *β*_0_ and {*β*_*l,h*_ }are regression intercept and coefficients, respectively. The classifier was trained with an *L*^1^-regularization penalty (weight 0.15) on attention features extracted from 20 experimentally resolved protein structures. Its strong generalization to previously unseen proteins demonstrated that ESM’s attention patterns encode residue–residue structural relationships, supporting the hypothesis of emergent structural information within large-scale protein language models.

## Methods

### Scoring metrics

Protein language models capture statistical regularities shaped by evolutionary constraints, and their likelihood estimates have been shown to correlate with functional fitness. For example, Hie et al. ^18^ demonstrated that viral evolution can be interpreted through a “grammar and context” framework, where mutations that preserve grammatical structure while altering antigenic context are more likely to promote immune escape. More broadly, Meier et al. ^19^ performed a large-scale evaluation across deep mutational scanning datasets and showed that several likelihood-based scores from masked language models can predict functional outcomes directly from sequence statistics.

Figure 1 provides an overview of the scoring metrics considered, which we describe in the following sections.

**Figure 1:**
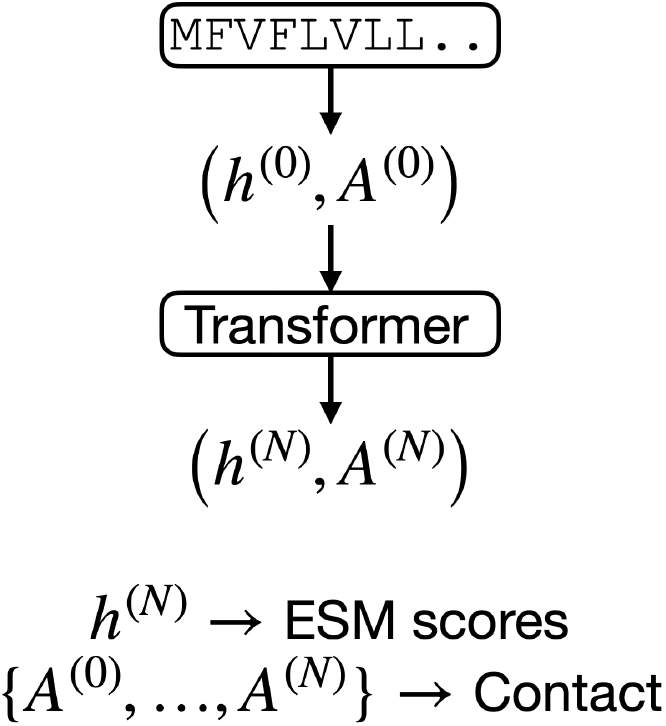
Schematic of scoring metrics.

### ESM scores

Motivated by these findings—and by the possibility that large structural deformations may accompany reduced fitness or functional disruption – we examine the three top-performing likelihood-based scoring methods identified in Meier et al. ^19^, each derived from conditional log-probabilities assigned by ESM.

#### Masked marginal

Let *x*^wt^ denote the wild-type sequence and *x*^mt^ the mutant sequence. For a mutation at position *i*, let *x*_*\i*_ denote the sequence with the residue at position *i* replaced by a mask token. The masked marginal score compares the log-probability of the mutant residue to that of the wild-type residue when the model conditions on the masked wild-type context:

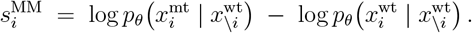

#### Wild-type marginal

The wild-type marginal conditions on the unmodified wild-type sequence evaluates how likely the mutant residue would be in the wild-type structural and evolutionary context:

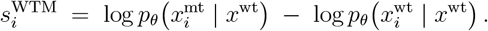

#### Mutant marginal

The mutant marginal conditions on the mutated sequence and compares how plausible the mutant residue is relative to the wild-type residue in the new context:

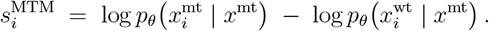

### Embedding distance

The last hidden representations 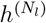 can be used for various downstream tasks. Here, down-stream tasks include unmasking for sequence generation with a regression head and structure prediction with folding blocks^14^. Therefore, 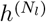 encodes rich information about the given protein.

We consider ℓ^1^ distance and cosine similarity (after flattening) between 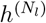 of the wild-type sequence and that of the mutated sequence.

### Contact difference

We aim to predict whether a single-point mutation induces a large structural deformation in the ESMFold-predicted tertiary structure, while avoiding the high computational cost of folding every mutant. The key idea is that ESM’s predicted residue–residue contact probabilities encode coarse-grained geometric information that can change detectably when a mutation substantially perturbs the structure.^2^

#### Contact-probability matrices

For an amino-acid sequence of length *L*, let

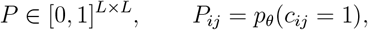

denote the matrix of predicted contact probabilities under a pretrained ESM model, where *c*_*ij*_ = 1 indicates that residues *i* and *j* are spatially proximal (e.g. within a threshold C_*α*_ distance). For a mutation at position *i* from its wild-type residue to *a* ∈ 𝒜, we write the corresponding contact matrix as

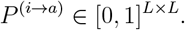

#### Local (row-wise) contact differences

A single-point mutation at position *i* most directly affects interactions involving residue *i*. We therefore consider the difference between the *i*-th rows:

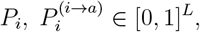

where *P*_*i*_ = (*P*_*i*1_, …, *P*_*iL*_) encodes the contact-probability distribution between residue *i* and all other residues. We define the local contact-difference vector

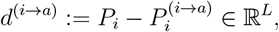

and measure its magnitude using standard vector norms, e.g.

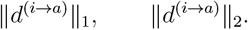

Large values of these norms indicate substantial changes in the predicted interaction pattern of residue *i*, suggesting a higher likelihood of local structural deformation in the folded mutant.

A canonical way to extend this approach to cases of multiple mutations is to extract the submatrix and then compare matrix norms.

#### Global (matrix-wise) contact differences

Although mutations primarily affect interactions involving residue *i*, they may also induce nonlocal changes in contacts among other residues. To account for this possibility, we consider the full matrix difference

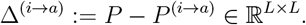

For completeness, we describe how each norm used in our analysis is computed. Let *M* = (*M*_*ij*_) ∈ ℝ^*L×L*^ be any matrix (in our application, *M* = Δ^(i→a)^).

#### Frobenius norm

The Frobenius norm measures the overall magnitude of the perturbation by summing the squared entrywise differences:

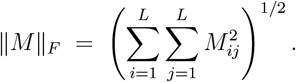

#### Entrywise ℓ_1_ norm

The entrywise ℓ_1_ (or 1, 1)-norm aggregates absolute differences across all pairs of residues:

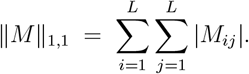

#### Induced ℓ_1_ operator norm

The operator 1-norm measures the maximum columnwise deviation:

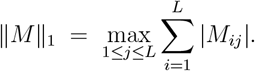

#### Induced ℓ_2_ operator norm (spectral norm)

The 2-operator norm corresponds to the largest singular value of *M*:

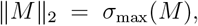

where *σ*_max_(*M*) denotes the largest singular value. This norm captures the largest linear amplification of the perturbation.

Together, the local and global contact-difference measures provide a computationally inexpensive surrogate for estimating structural deformation without performing full structure prediction for all mutants.

## Results

### Correlation study

To quantify structural changes between wild-type and mutant models, we use two metrics: root-mean-square deviation (RMSD) and strain^8^. RMSD is computed as the ℓ^2^ distance between the aligned vectors of C_*α*_ coordinates, normalized by 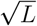 for a protein of length *L*. Strain provides a complementary, residue-localized measure of deformation. For each position *i*, the strain is defined as

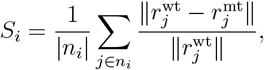

where *n*_*i*_ denotes the set of residues in spatial proximity to position *i*, and *r*_*j*_ is the *C*_*α*_ coordinate of residue *j*. A global strain score is then obtained by averaging over all residues, 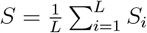 For additional computational details for strain, including structural alignment and preprocessing steps, we refer the reader to PDAnalysis.^3^ For the protein language model, we used ESM2 with 650 million parameters.

#### SARS-CoV-2 spike protein

Table 1 reports Spearman rank correlations between backbone RMSD (and strain, separately) and a collection of contact-map perturbation metrics and sequence-based scores for the SARS-CoV-2 spike protein.^4^ All contact-based metrics show positive and highly significant correlations, indicating that larger perturbations of the predicted contact map are consistently associated with greater structural displacement in the ESMFold output. However, their relative performance differs across metric families.

**Table 1:** Study results on SARS-CoV-2 spike protein (200 random single mutations). Overall, embedding distance performed well.

| category | method | correlation w/ RMSD | correlation w/ strain |
| --- | --- | --- | --- |
| ESM scores | masked marginal | -0.452 (1.8e-11) | -0.375 (4.4e-08) |
|  | wildtype marginal | **-0.507** (1.8e-14) | -0.449 (2.6e-11) |
|  | mutant marginal | -0.249 (3.8e-04) | -0.161 (2.2e-02) |
| Embeddings | embedding distance | **0.552** (2.4e-17) | **0.616** (2.9e-22) |
|  | cosine similarity | -0.449 (2.6e-11) | **-0.535** (3.5e-16) |
| Contact | $\ell_1$ | 0.280 (6.0e-05) | 0.360 (1.6e-07) |
| | $\ell_2$ | 0.272 (9.9e-05) | 0.332 (1.6e-06) |
| | operator $\ell_1$ | 0.241 (5.7e-04) | 0.332 (1.6e-06) |
| | operator $\ell_2$ | 0.258 (2.2e-04) | 0.336 (1.1e-06) |
|  | Frobenius 1 | 0.439 (7.6e-11) | 0.500 (4.9e-14) |
|  | Frobenius 2 | 0.337 (1.0e-06) | 0.394 (7.5e-09) |

Among contact map-based scores, the operator norms induced by vector (ℓ^1^ and ℓ^2^) produce the weakest correlations. This well matches our intuition, since induced operator norms are meaningful when matrices can be interpreted as linear transformations between two spaces; contact map matrices lack such an interpretation. Vector-based metrics still performed worse than element-wise matrix norms. Because these metrics compress the contact-map difference into row- or columnwise aggregates, they discard most residue–residue information; their lower correlations suggest that structural deformation is not well summarized by such coarse projections. The strongest performance is achieved by the elementwise Frobenius norms (`frobenius_1` and `frobenius_2`), which retain full residue-pair resolution. Their leading correlations with both RMSD and strain suggest that distributed, fine-grained contact-map changes offer the most informative signal for predicting geometric deviation.

The ESM sequence-based scores (masked marginal, wild-type marginal, and mutant marginal) display significant *negative* correlations with RMSD and strain. Mutations predicted by ESM to have lower marginal likelihoods – i.e., to be less evolutionarily plausible – tend to produce larger structural deformations in the ESMFold predictions. This pattern is consistent with the expectation that mutations strongly perturbing the native fold are typically deleterious and therefore assigned a lower probability by the language model.

Embedding distance attains the strongest correlations among all features, outperforming both contact-map metrics and sequence-based marginal scores. Hie et al. ^18^ previously combined embedding distance with marginal probabilities to model viral evolution and reported strong agreement with experimentally measured antibody escape. Their approach favors mutations that simultaneously move far in embedding space and retain high evolutionary plausibility. Because marginal probabilities often correlate with fitness^19^ – measured by binding affinity with ACE2 in this context – the embedding-distance term may amplify changes to epitope-paratope interactions. Our finding that embedding distance alone is strongly associated with ESMFold-predicted deformation is broadly consistent with this interpretation. However, more detailed analysis is warranted, as ESMFold’s structure module directly consumes hidden representations from the same language model, which may partly account for the high correspondence.

#### SARS-CoV-2 spike protein (AF2, multi-mutant variants)

To evaluate whether our scoring metrics remain informative when multiple substitutions occur simultaneously, we analyzed AF2-predicted structures of SARS-CoV-2 spike variants containing five mutations per sequence. Unlike the single-mutant datasets considered previously, these variants contain 5 mutations, requiring us to extract the corresponding submatrices of the predicted contact maps before applying vector- and matrix-based norms. As shown in Table 2, marginal-likelihood scores maintain consistent negative correlations with RMSD and strain, and embedding distance still produces the strongest positive correlation with strain.

**Table 2:** Correlation study results for AF2-predicted SARS-CoV-2 spike variants containing five substitutions per sequence. Correlations are substantially weaker than in single-mutant datasets, likely due in part to these variants lying far outside the evolutionary distribution modeled by ESM (average masked-marginal score (≈ − 9.9). Marginal scores nonetheless show consistent negative associations with structural deviation, and embedding distance yields the strongest positive correlation with strain.

| category | method | correlation w/ RMSD | correlation w/ strain |
| --- | --- | --- | --- |
| ESM scores | masked marginal | **-0.129** (6.8e-02) | **-0.158** (2.5e-02) |
|  | wildtype marginal | -0.079 (2.7e-01) | -0.153 (3.1e-02) |
|  | mutant marginal | **-0.126** (7.6e-02) | -0.133 (6.0e-02) |
| Embeddings | embedding distance | 0.114 (1.1e-01) | **0.182** (9.9e-03) |
|  | cosine similarity | -0.022 (7.6e-01) | -0.119 (9.2e-02) |
| Contacts | $\ell_1$ | -0.041 (5.7e-01) | 0.016 (8.2e-01) |
| | $\ell_2$ | -0.054 (4.5e-01) | 0.000 (1.0e+00) |
| | operator $\ell_1$ | -0.041 (5.7e-01) | -0.033 (6.4e-01) |
| | operator $\ell_2$ | -0.026 (7.2e-01) | 0.029 (6.8e-01) |
|  | Frobenius 1 | 0.074 (3.0e-01) | 0.122 (8.6e-02) |
|  | Frobenius 2 | 0.014 (8.4e-01) | 0.072 (3.1e-01) |

But, correlations in this multi-mutant setting are substantially weaker than in the single-mutation experiments. One likely contributing factor is that these variants are far from the evolutionary manifold captured by the language model. The average masked marginal score across the five mutated positions is approximately − 9.9, indicating that the model assigns low probability to these sequences. Figure 2 presents histograms of marginal scores for three different datasets. The p-value for two sample t-test for the first two datasets was 3.0e-42, suggesting a statistically significant difference between the two masked marginal score distributions. Such sequences are expected to be unlikely in nature, which may distort the performance of the structure prediction model and therefore weaken the correlation. To support this claim, we restricted the analysis to those mutants that have a masked marginal score of more than −7. The rank correlation between the embedding distance and the strain was increased to 0.307. The rank correlation between the embedding distance and the rmsd was increased to 0.158, but remained statistically insignificant (p=1.6e-1).

**Figure 2:**
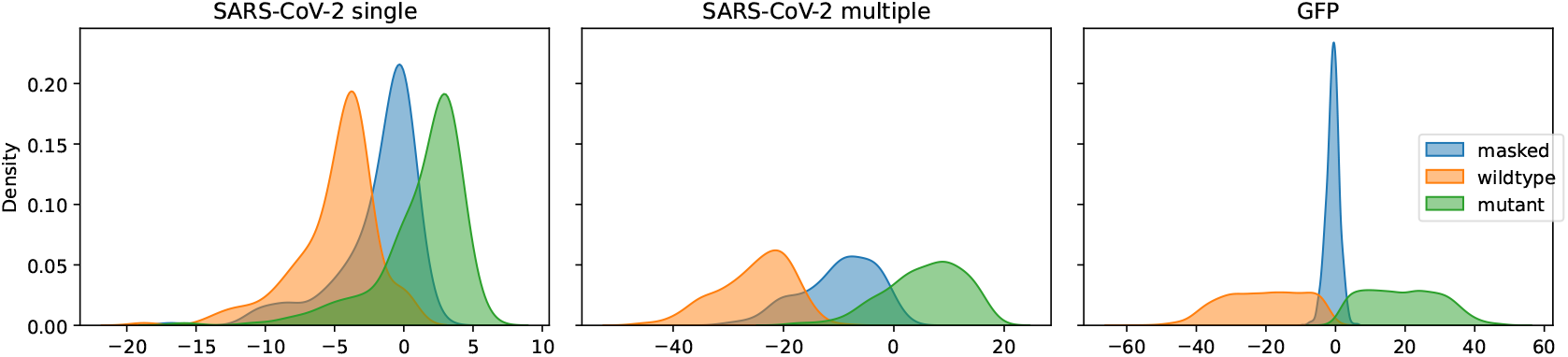
Masked marginal scores for 1) 200 random single mutations for SARS-CoV-2 spike protein, 2) 200 random 5 mutations for SARS-CoV-2 spike protein, and 3) 2,312 natural mutations for green fluorescent protein.

#### Green fluorescent protein

We further evaluated our scoring metrics on a large mutational dataset of green fluorescent protein (GFP), consisting of 2,312 mutants originally generated to study the GFP fitness landscape^20^. McBride et al. ^8^ generated AF2 predictions for this dataset, to study whether AF2 captures mutational effects or not. Unlike synthetically generated multiple mutants for the SARS-CoV-2 spike protein, this dataset consists of synthesizable proteins. Figure 2 presents masked marginal scores’ distributional differences between synthetically generated mutations and natural mutations (p-value *<* 10^−324^ for SARS-CoV-2 Multiple vs. GFP). As in the SARS-CoV-2 spike protein analysis, we extracted contact-map submatrices corresponding to the mutated positions before computing vector- and matrix-based perturbation metrics. As shown in Table 3, the wild-type and mutant marginal scores exhibit the strongest correlations overall, with the wild-type marginal negatively correlated with RMSD and strain (*ρ* = −0.583 and *ρ* = − 0.712), while the mutant marginal shows stronger positive correlations (*ρ* = 0.603 and *ρ* = 0.726). This sign reversal relative to single-mutation datasets reflects the altered statistical context introduced by multiple substitutions and suggests that the mutant marginal score may be unreliable for *in silico* screening. Among contact-based metrics, the entrywise ℓ^1^ and ℓ^2^ norms for submatrices achieve the highest correlations, consistent with their capacity to capture distributed perturbations across multiple mutated sites. Notably, embedding distance continues to show strong performance (*ρ* = − 0.538 and ρ = 0.640), demonstrating that global changes in ESM representations remain predictive of AF2-derived structural deformation even in the presence of complex, multi-residue mutations.

**Table 3:**
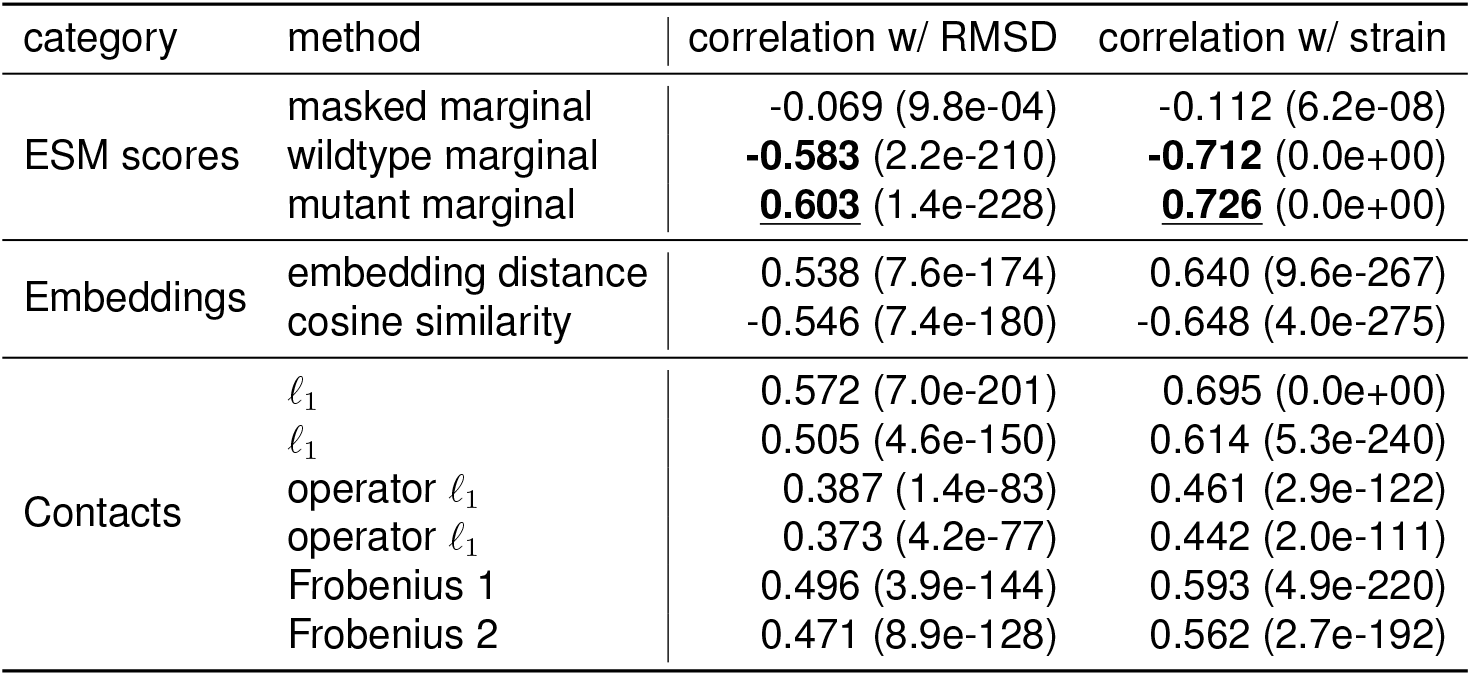
Study results for green fluorescent protein (GFP). Wild-type and mutant marginal scores show the strongest correlations with RMSD and strain, though the mutant marginal exhibits an opposite sign compared to single-mutation datasets, limiting its utility for *in silico* screening. Among contact-based metrics, the entrywise ℓ^1^ and ℓ^2^ norms perform best. Because GFP variants contain multiple mutations, ℓ-metrics were computed by extracting and vectorizing the corresponding contact-map submatrices. Embedding distance also shows robust correlations with AF2-predicted structural deformation.

| category | method | correlation w/ RMSD | correlation w/ strain |
| --- | --- | --- | --- |
| ESM scores | masked marginal | -0.069 (9.8e-04) | -0.112 (6.2e-08) |
|  | wildtype marginal | **-0.583** (2.2e-210) | **-0.712** (0.0e+00) |
|  | mutant marginal | **0.603** (1.4e-228) | **0.726** (0.0e+00) |
| Embeddings | embedding distance | 0.538 (7.6e-174) | 0.640 (9.6e-267) |
|  | cosine similarity | -0.546 (7.4e-180) | -0.648 (4.0e-275) |
| Contacts | $\ell_1$ | 0.572 (7.0e-201) | 0.695 (0.0e+00) |
| | $\ell_1$ | 0.505 (4.6e-150) | 0.614 (5.3e-240) |
| | operator $\ell_1$ | 0.387 (1.4e-83) | 0.461 (2.9e-122) |
| | operator $\ell_1$ | 0.373 (4.2e-77) | 0.442 (2.0e-111) |
|  | Frobenius 1 | 0.496 (3.9e-144) | 0.593 (4.9e-220) |
|  | Frobenius 2 | 0.471 (8.9e-128) | 0.562 (2.7e-192) |

#### Summary of scoring performance and choice of method

Across all datasets, embedding distance emerged as the most consistent and reliable predictor of structural deformation. In the SARS-CoV-2 spike single-mutation study (ESMFold), it achieved the highest correlations with both RMSD and strain, outperforming all contact-based measures and ESM marginal scores. In the more challenging SARS-CoV-2 spike setting with five simultaneous mutations (AF2), embedding distance remained the strongest positive correlate with strain, whereas contact-based metrics degraded substantially and marginal scores weakened. In the GFP multi-mutant dataset (AF2), embedding distance again exhibited high correlations, matching or surpassing the best-performing contact-map metrics. Taken together, these results show that embedding distance offers the most robust and generalizable signal of structural perturbation across proteins, mutation regimes, and structure-prediction backbones. We therefore adopt embedding distance as the primary scoring method for the subsequent application.

### High-throughput screening

Rift Valley fever virus (RVFV) causes a mosquito-borne zoonotic disease. We applied our method to estimate mutational effects on structural changes in the MP-12 strain, an effective vaccine strain. Our analysis targeted the M-segment,^5^ which is central to antigenic function and contains two attenuating mutations relative to the ZH501 strain^21^.

A single ESMFold prediction for the 1197-residue sequence requires roughly 85 seconds on an NVIDIA H200 GPU. Exhaustively evaluating all 22,724 (excluding position 1) single mutants would therefore require more than 22 days, making full-coverage structure prediction computationally prohibitive.^6^

Without performing any structure prediction, we computed embedding distances between the wild-type sequence and all single mutants, which only took 23 minutes in the same computing environment. Based solely on these scores, we selected the top 100 and bottom 100 mutations – corresponding to the largest and smallest predicted embedding shifts, respectively. We then ran ESMFold on this reduced set of 200 mutations to assess the effectiveness of the screening strategy.

Figure 3 shows screening results for RVFV using embedding distance. Mutations in the top-100 group produced markedly larger structural deviations, with a mean RMSD of **12.5** (strain: **0.123**). In contrast, the bottom-100 group exhibited substantially smaller perturbations, with a mean RMSD of **3.16** (strain: **0.0590**) **(a)**. This pronounced separation demonstrates that embedding distance serves as an effective and reliable prescreening metric for identifying mutations likely to cause significant structural rearrangements, enabling orders-of-magnitude reductions in required ESMFold inference. Panel **(b)** shows representative ESMFold-predicted structures visualized using UCSF ChimeraX^22^. For visual clarity, we display only residues 450–590, which include both mutated positions. The clear structural deviation of the 579W mutant (blue) relative to the wild type (red) illustrates how large embedding-distance shifts correspond to substantial local and quasi-local structural rearrangements.

**Figure 3:**
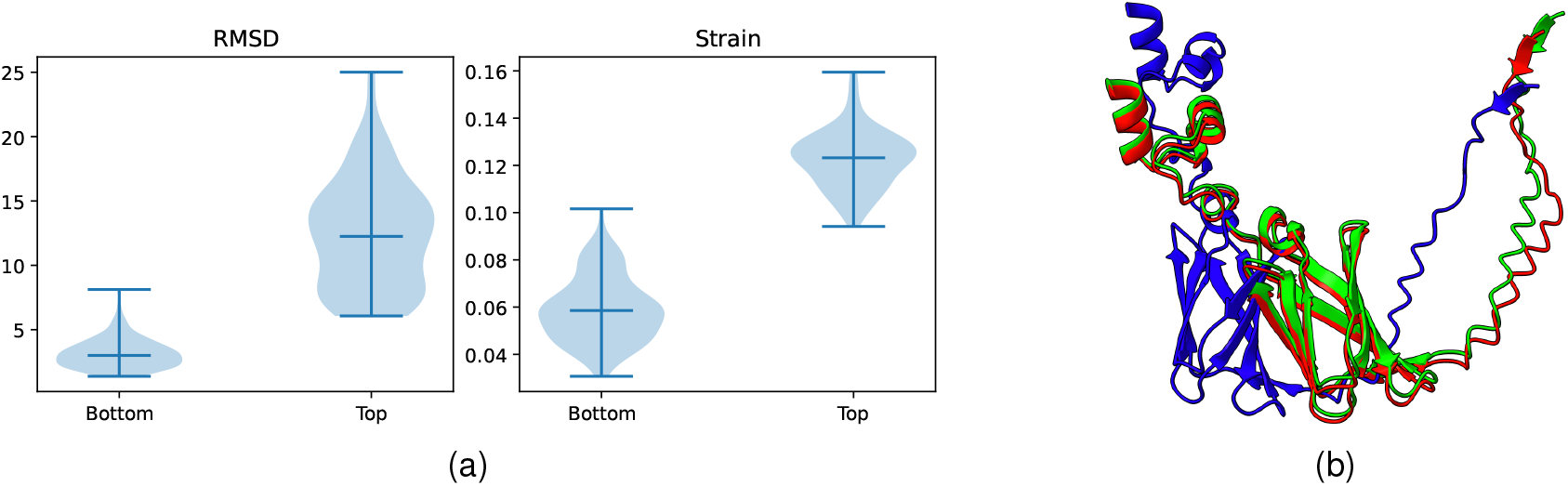
Screening results for RVFV using embedding distance. **(a)** The distributions of RMSD and strain for the top- and bottom-scoring mutation groups selected by embedding distance. Differences between groups are statistically significant (two-sample t-test: RMSD *p* = 2.0 *×* 10^−52^; strain p = 5.9 × 10^−85^). **(b)** Visualization of ESMFold-predicted RVFV M-segment structures (residues 450–590). Colors indicate wild type (red), the lowest-embedding-distance mutant 462I (green), and the highest-embedding-distance mutant 579W (blue). The pronounced deviation of 579W highlights the correspondence between large embedding-distance shifts and substantial structural perturbation.

## Discussion

### Conclusion

In this work, we introduced a computationally efficient approach for predicting whether missense mutations are likely to induce substantial structural deformation in ESMFold-predicted protein models. By leveraging emergent structural signals encoded in ESM – either through hidden-representation changes or through differences in predicted residue–residue contact probabilities – we demonstrated the possibility of extracting several sequence-derived scores that correlate with RMSD as well as strain between wild-type and mutant structures. Among these, ESM embedding distance consistently demonstrated strong performance across multiple proteins, enabling the rapid identification of structure-disrupting mutations without requiring full structure prediction for every variant, whose computational cost is prohibitively high. This ability to efficiently prescreen large mutational spaces provides a scalable and practical tool for high-throughput protein engineering, viral antigen analysis, and early-stage computational design workflows.

### Limitations

Despite promising results, several limitations remain. First, the correlations observed in single-mutant settings weaken substantially for variants containing multiple substitutions. Our analysis suggests that this degradation is linked to mutations driving sequences farther from the evolutionary manifold captured by the language model, as indicated by strongly negative marginal-likelihood scores. In such cases, ESM embeddings and contact predictions may no longer provide reliable structural signals. Second, our evaluation relies on ESMFold and AF2-generated structures, which may contain prediction artifacts for destabilizing or out-of-distribution sequences. Finally, we have focused on correlation-based evaluation rather than direct prediction of deformation magnitudes or stability changes, which may require more specialized models.

### Future directions

Several avenues could enhance the utility and generality of our approach. One direction is to combine complementary ESM-derived scores – such as embedding distances, marginal likelihoods, and contact-map perturbations – into a unified predictor, potentially via a lightweight regression or ranking model. Another potential direction is to re-fit or fine-tune the contact-prediction head on structurally relevant families or phylogenetically related proteins, which may sharpen the correspondence between contact differences and structural deformation. Evo-tuning^23^ or domain-specific representation learning could further improve performance for proteins with rich evolutionary context. Extending our analysis to experimentally determined mutant structures would help validate the robustness of our findings beyond model-generated data. Finally, the proposed scoring framework naturally integrates into multi-objective optimization pipelines^24^, enabling sequence design strategies that jointly consider fitness, function, and structural stability. We leave these investigations for future work.

## Acknowledgments

This work has been supported by the ARPA-H Award #1AY1AX000053-01.

## Data availability

The data used in this article are publicly available, and the relevant information needed for accessing the data can be found in the article itself.

## Footnotes

1 after symmetrized and APC-corrected.

2 Because ESMFold does not require multiple sequence alignments (MSAs), inference is faster than AlphaFold. Nonetheless, folding all 19*L* single-point mutants remains costly for long proteins.

3 https://github.com/mirabdi/PDAnalysis

4 Uniprot id: P0DTC2

5 The amino acid sequence can be found at GenBank accession ABD38821.1.

6 Runtime could be reduced with optimized batching, but such engineering considerations lie outside the scope of this work.

## Notes

### Competing Interest Statement

The authors have declared no competing interest.

